# Multiple hidden processes complicate phylogenomic inference of deep Basidiomycota relationships

**DOI:** 10.1101/170696

**Authors:** Arun N. Prasanna, Daniel Gerber, Kijpornyongpan Teeratas, M. Catherina Aime, Vinson Doyle, Laszlo G. Nagy

## Abstract

Resolving deep divergences in the fungal tree of life remains a challenging task even for analyses of genome-scale phylogenetic datasets. Relationships between Basidiomycota subphyla, the rusts (Pucciniomycotina), smuts (Ustilaginomycotina) and mushroom forming fungi (Agaricomycotina) represent a particularly challenging situation that posed problems to both traditional multigene and genome-scale phylogenetic studies. Here, we address basal Basidiomycota relationships using three different phylogenomic datasets, concatenated and gene tree-based analyses and examine the contribution of several potential sources of uncertainty, including fast-evolving sites, putative long-branch taxa, model violation and missing data. We inferred conflicting results with different datasets and under different models. Fast-evolving sites and oversimplified models of amino acid substitution favored the grouping of smuts with mushroom-forming fungi, often leading to maximal bootstrap support in both concatenation and Astral analyses. The most conserved datasets grouped rusts with mushroom forming fungi, although this relationship proved labile, sensitive to model choice, different data subsets and missing data. Excluding putative long branch taxa, genes with the highest proportions of missing data and/or genes with strong signal failed to reveal a consistent trend toward one or the other topology, suggesting that additional sources of conflict are at play too. Our analyses suggest that topologies uniting smuts with mushroom forming fungi can arise as a result of inappropriate modeling of amino acid sites that might be prone to systematic bias. While concatenated analyses yielded strong but conflicting support, individual gene trees mostly provided poor support for rusts, smuts and mushroom-forming fungi, suggesting that the true Basidiomycota tree might be in a part of the tree space that is difficult to access using both concatenation and gene tree based approaches. Thus, basal Basidiomycota relationships remain unresolved and might represent a phylogenetic problem that remains contentious even in the genomic era.

## Introduction

Accurately resolving phylogenetic relationships across the Tree of Life has been one of the central challenges for evolutionary biologists. Phylogenetic datasets relying on one to a few genes have left many phylogenetic questions open, for which the use of genomic data appeared to be a magic bullet(Delsuc, et al. 2005; Gee 2003; Misof, et al. 2014; Philippe, et al. 2005; Prum, et al. 2015; Rokas, et al. 2003). Genome-scale datasets are 10x - 1000x the size of traditional phylogenetic datasets, virtually eliminating data availability as the limiting factor of resolving phylogenetic questions, a great promise for several historically recalcitrant nodes(Misof, et al. 2014; Niehuis, et al. 2012; Rokas, et al. 2003; Telford, et al. 2015). Despite the early promise of initial genome-scale studies, phylogenomic datasets present fundamentally new challenges in terms of dataset assembly, effects of missing data and the strength and sources of incongruence(Galtier and Daubin 2008; Jeffroy, et al. 2006; Kumar, et al. 2012; Philippe, et al. 2011; Philippe, et al. 2005; Soltis, et al. 2004).

Compared to traditional phylogenetics, phylogenomic datasets often yield highly resolved, often maximally supported trees. Increasing the amount of data alleviates some sources of incongruence, but not others: inadequate modeling of the data can lead to systematic biases in the estimation procedure(Jeffroy, et al. 2006; Kumar, et al. 2012; Philippe, et al. 2011; Philippe, et al. 2005). In such cases, there is no guarantee that the true phylogenetic signal dominates the inferred topology, even if it is strongly supported, as systematic errors can inflate support values for incorrect topologies under some circumstances(Parks, et al. 2012; Philippe, et al. 2011; Philippe, et al. 2005; Phillips, et al. 2004; Rodriguez-Ezpeleta, et al. 2007; Sharma, et al. 2014). The source of systematic errors can be several biological (e.g. heterotachy, compositional bias, rate heterogeneity, incomplete lineage sorting, HGT, etc…)(Hallstrom and Janke 2010; Li, et al. 2016; Parks, et al. 2012; Phillips, et al. 2004; Rodriguez-Ezpeleta, et al. 2007; Romiguier, et al. 2016; Sharma, et al. 2014; Sharma, et al. 2015; Smith, et al. 2015; Szollosi, et al. 2012; Whelan, et al. 2015) or technical issue (e.g. biased taxon or gene sampling, orthology inference, poor model fit)(Chen, et al. 2015; Dell’Ampio, et al. 2014; Dunn, et al. 2008; Hejnol, et al. 2009; Hosner, et al. 2016; Lanfear, et al. 2014; Misof, et al. 2013; Pisani, et al. 2015; Simon, et al. 2012; Smith, et al. 2015; Whelan, et al. 2015; Yang and Smith 2014); these can confound phylogenomic analyses and lead to biased or incorrect trees(Kumar, et al. 2012; Philippe, et al. 2011).

Fungi have been at the forefront of phylogenomics due to the relative ease of sequencing their genomes(Grigoriev, et al. 2014), and have been among the first groups to be subjected to phylogenomic analyses(Aguileta, et al. 2008; Fitzpatrick, et al. 2006; Robbertse, et al. 2006; Rokas, et al. 2003), yet, several deep divergences in the fungal tree await resolution. Basal relationships of the Basidiomycota have been particularly recalcitrant (Aime, et al. 2006; Hibbett 2006; Hibbett, et al. 2007; Kohler, et al. 2015; Matheny, et al. 2007; Nagy, et al. 2016; Padamsee, et al.) both and recent phylogenomic treatments have recovered strongly supported contradicting relationships. Ultrastructural characters can be difficult too to interpret (Lutzoni, et al. 2004) but seem to favor the grouping of smuts with mushroom-forming fungi, based on the development of membrane-bounded septal pores, and swollen margins in both. The divergence of basal basidiomycete clades was estimated at ca 521 MYR(Floudas, et al. 2012) and comprises three large clades, the rusts (Pucciniomycotina), the smuts (Ustilaginomycotina) and the mushroom-forming fungi (Agaricomycotina). Traditional multilocus studies have generally found weak support for various groupings of these clades(Aime, et al. 2006; Berres, et al. 1995; Hibbett, et al. 2007; James, et al. 2006; Matheny, et al. 2007; Swann and Taylor 1993), whereas genome-scale studies have provided often strongly supported but conflicting relationships. In most previous studies smuts and mushroom-forming fungi grouped together to the exclusion of rusts (referred to as R(S,M) topology hereafter) (Ebersberger, et al. 2012; Kohler, et al. 2015; Padamsee, et al.; Sharma, et al. 2015; Zajc, et al. 2013), although rusts as the sister group of smuts (hereafter: M(R,S) topology) (Kohler, et al. 2015) and rusts as the sister group of mushroom-forming fungi (hereafter: S(R,M) topology) (Medina, et al. 2011; Nagy, et al. 2014; Riley, et al. 2014) have also been reported. Further, analyses of the same datasets under different models or methods have yielded contradicting results(Kohler, et al. 2015), suggestive of evolutionary processes not captured by the models or methods used. For example, taxon sampling was biased towards mushroom-forming fungi in most previous phylogenomic studies, with only 1-2 rust and smut species analyzed. Biased taxon sampling can compromise phylogenomic inference(Dunn, et al. 2008; Philippe, et al. 2011; Pisani, et al. 2015; Simon, et al. 2012) and has been suggested as a potential factor underlying the difficulty of resolving basal Basidiomycota relationships(Nagy, et al. 2016). Notably, both rusts and smuts have undergone massive gene losses compared to mushroom-forming fungi and Ascomycota (Nagy, et al.2014), which may impact the number of genes available for phylogenetic inference and thus the reconstructed relationships. It has also been suggested that basal relationships of the Basidiomycota might have been shaped by fast successive speciation events and thus be better described as a hard polytomy (Kohler, et al. 2015).

The selection of orthologous groups of genes is one of the most important determinants of dataset quality (Dunn, et al. 2008; Simon, et al. 2012; Smith, et al. 2015; Whelan, et al. 2015; Yang and Smith 2014). Unwanted inclusion of non-orthologous genes (e.g. deep paralogs, horizontally acquired genes) can introduce both noise and incongruence in the datasets, whereas the inclusion of too much (non-random) missing data in concatenated analyses can influence phylogenetic signal (Chen, et al. 2015; Dell’Ampio, et al. 2014; Hejnol, et al. 2009; Hosner, et al. 2016; Misof, et al. 2013; Streicher, et al. 2016; Xi, et al. 2016). It is widely accepted that tree-based methods are most accurate for determining ortholog/paralog relationships (Boussau, et al. 2013; Gabaldon 2008; Kocot, et al. 2013; Kocot, et al. 2017; Kristensen, et al. 2011; Yang and Smith 2014), however, they have surprisingly rarely been the choice in phylogenomic dataset assembly in favor of simpler, blast- or similarity-based methods (for exceptions see (Dunn, et al. 2008; Hejnol, et al. 2009; Smith, et al. 2011).

In this study we set out to identify the causes of the difficulties in resolving the deepest basidiomycete nodes and to infer a robust phylogenetic hypothesis for the early evolution of these. We significantly increase the number of sampled rust and smut species, examine the effects of gene and site selection, model choice and signal-to-noise ratios. We employ an gene tree-based orthogroup inference pipeline that minimizes the chance for including non-orthologous genes and thus technical sources of gene tree conflict. While dissecting the sources of uncertainty in the historically recalcitrant node at the base of Basidiomycota, we detect multiple sources of bias in the datasets that lead to persistent challenges for resolving ancient Basidiomycete relationships.

## Materials and Methods

### Taxon sampling

We used complete genome sequences of 67 species, representing the three major clades of the Basidiomycota plus 10 Ascomycetes and *Phycomyces*, *Mucor* and *Rhizophagus* as outgroups (Table S1). The Basidiomycota included 32 Agaricomycotina (mushroom-forming fungi, including Wallemiomycetes), 12 species of rusts (Pucciniomycotina) and 10 species of smuts (Ustilaginomycotina).

### Blast, clustering and the selection of orthologous groups

We performed all-vs-all blast on non-redundant predicted protein sequences in the 67 input genomes using mpiblast 1.6.0. Proteins showing significant hits were clustered based on similarity using the Markov clustering algorithm implemented in Mcl 14-137(van Dongen 2000) with an inflation parameter of 2.0. Clusters containing 33 to 134 proteins (50 to 200% of the species analyzed) were further analyzed as these are most likely to contain conserved, singlecopy genes for the highest number of species. Multiple alignments and gene trees were estimated for each of these clusters using PRANK v.140603 (Loytynoja and Goldman 2008) and RAxML 7.2.3 (Stamatakis 2014), respectively (both with default parameters). The WAG model of protein sequence evolution with gamma-distributed rate heterogeneity was used in RAxML. We then screened the resulting gene family trees for gene duplications and retained only those that had no duplications at all (only a single protein per species) or terminal duplications only (i.e. inparalogs), using a custom perl script (available from the authors upon request). Clusters containing deep paralogs were discarded. Of inparalogs we retained the one with the shortest root-to-tip patristic distance. From the resulting gene families we further excluded ones that contained overly divergent, potentially non-orthologous proteins by calculating the contribution of each terminal branch length to the total tree length; gene families in which a single gene accounting for >60% of the total tree length were eliminated (dos Reis, et al. 2012).

### Data filtering, concatenation and phylogenetic analyses

We used GBlocks 0.91b (Castresana 2000) to remove poorly aligned regions from the alignments using three levels of stringency: first, with default parameters, second with ‘−b2=50 − b3=10 −b4=5’ and third with ‘−b2=40 −b3=20 −b4=2’. For comparison, we also analyzed the original gene family alignments (no sites removed). Next, we concatenated gene families that had representatives in >50% of the species and a final trimmed length >50 amino acid sites. The resulting four datasets were analyzed using the PTHREADS version of RAxML 7.2.3 (Stamatakis 2014), PhyloBayes 4.1 (Lartillot, et al. 2013) and MrBayes 3.2.6 (Ronquist and Huelsenbeck 2003). For maximum likelihood and MrBayes analysis we partitioned the dataset into gene families and used the best-fit model for each partition with gamma-distributed rate heterogeneity. Bootstrap analyses were run in 100 replicates. The GBlocks-untreated dataset was not subjected to bootstrapping due to the prohibitive run time of the analysis. PhyloBayes was run with 2 independent replicates with one chain each, for 8000 cycles under the CAT-GTR model of sequence evolution. The first 1000 cycles were discarded as burn-in. MrBayes was run with 2 independent replicates with 4 chains each, for 1,000,000 generations and a partitioned model (WAG for each partition) with 250,000 generations as burn-in. Convergence was assessed by inspecting clade posterior probabilities and by the bpcomp and tracecomp functions of Phylobayes.

To examine the information content of the datasets with regard to the branching of smuts, rusts and mushroom-forming fungi, we performed four-quartet likelihood mapping (Strimmer and von Haeseler 1997) in Tree-puzzle (Schmidt, et al. 2002). This was done under the WAG model with gamma-distributed rate heterogeneity (4 categories) and examined 100,000 quartets. To address saturation levels of the datasets, we calculated uncorrected and patristic distances from the trimmed single-gene alignments and corresponding ML gene trees using the ‘dist.hamming’ and ‘cophenetic.phylo’ functions of the phangorn 2.1.1 (Schliep 2011) and ape 4.0 (Paradis, et al. 2004) R packages, respectively.

### Phylogeny based on gene content

We assessed support for each of the three alternative conformations of rusts, smuts and mushroom-forming fungi based on gene presence/absence data in the 67 fungal genomes. A binary presence-absence (PA) matrix was constructed for each of the protein clusters (50,971) against all the species (67). No distinction was made in the coding between single-copy and multicopy genes. Next, a maximum likelihood tree was inferred using RAxML 7.2.3 with a GAMMA model of rate heterogeneity and 1000 bootstrap replicates.

We calculated the number of gene family origins and losses needed to explain the data under the two competing conformations as an independent test of topologies. To this end, we manually rearranged the ML tree in Mesquite 4 (Maddison and Maddison 2009) to create trees with two alternative topologies: Rusts + Agaricomycetes and Smuts + Agaricomycetes. We then mapped gene family gains and losses using Dollo parsimony, recording the total number of gains and losses. The two competing topologies were then compared using the approximately unbiased test as implemented in CONSEL (Shimodaira and Hasewaga 2001).

### Coalescent-based analyses

To account for gene tree incongruence (e.g. resulting from incomplete lineage sorting) we used the Accurate Species Tree Algorithm (ASTRAL-II), to synthesize species trees from individual gene trees (Mirarab, et al. 2016; Mirarab and Warnow 2015), rather than concatenated data. We inferred gene trees from each of the trimmed gene family alignments for the 314, 824 and 901 gene datasets in RAxML under the WAG+G model with 100 bootstrap replicates. Astral-II 4.7.8 was used to summarize gene trees into a species tree with 100 multi-locus bootstrap replicates.

### Removal of fast-evolving amino acid sites

We used TIGER 1.02 (Cummins and McInerney 2011) to sort amino acid sites into 20 rate categories. Next, we sequentially removed the fastest evolving sites from each concatenated dataset and inferred ML trees with 100 bootstraps (as above) for each dataset for the 901G, 824G and 314G datasets. We removed additional rate categories until Rusts, Smuts, Mushroom-forming fungi and Ascomycetes were not monophyletic, which presumably marked the loss of a significant portion of the phylogenetic signal in the dataset.

### Analyses of hard polytomies

We tested the hypothesis that the Pucciniomycotina, Ustilaginomycotina and Agaricomycotina diverged from each other in fast successive speciation events - and thus, their relationships would be best described by multifurcation. Phycas 2.2.0 (Lewis, et al. 2010) was used to run Bayesian MCMC analyses that allow polytomic trees. We ran 100,000 cycles with 25,000 cycles as burn-in and an unpartitioned GTR+G model of sequence evolution on the 314G dataset (our attempts to run partitioned analyses failed due to the prohibitive memory requirements of the analyses). We ran analyses under a range of flat to polytomy-friendly priors, including polytomy and resolution class priors with setting the mcmc.topo_prior_c parameter between 1 and 3. Analyses were run in triplicates.

## Results

### Dataset completeness, decisivity

Clustering at an inflation parameter of 2.0 yielded 5,257 clusters that contained 33 to 134 proteins (50-200% of the number of species), of which 1,327 clusters contained a single gene per species or inparalogs only. Three alignments containing overly divergent sequences (contributing >60% of total tree length) were excluded from further analyses. Of the remaining clusters, 950 contained more than 33 species, corresponding to at least half of the species in the analyses. We removed ambiguously aligned regions from the alignments using three stringency levels in GBlocks, and retained the trimmed alignments if they contained >50 amino acid characters. This resulted in three sets of trimmed alignments comprising 314, 824 and 901 genes, which we designate as the 314G, 824G and 901G datasets. Further, we concatenated the 950 untrimmed alignments, which we hereafter refer to as the 950G dataset. The concatenated 314G, 824G, 901G and 950G datasets comprised 46,573, 165,465, 241,004 and 704,775 amino acid characters, respectively. Overall, the 314G dataset was the most conserved (mean pairwise ML distance 0.406) followed by the 824G (0.537), 901G (0.616) and 950G (0.962) datasets. Taxon occupancy was even across datasets (82% to 86%) with the 314G dataset having the fewest missing genes (14%), followed by the 824G, 901G and 950G datasets close to each other (all with 18% of the genes missing) (Fig S1). Mushroom-forming fungi were best represented in all datasets, with 89% - 92% of the orthologs present per species, followed by rusts (78% - 83%), smuts (74% - 82%) and Ascomycota (70% - 77%).

Most of the orthogroups in the three datasets had representatives in the rusts, smuts and mushroom-forming fungi, i.e. they were decisive sensu (Dell’Ampio, et al. 2014). Smuts have the highest number of orthogroups that had no representative at all (65-91), followed by rusts (11-14), whereas mushroom-forming fungi had representatives in all of the included orthogroups. Of the four datasets, the 314G dataset is the most conserved on average and is the least saturated, followed by the 824G, 901G and 950G datasets (Fig S2).

Phylogenetic signal conferred by individual gene families was examined by ML bootstrapping of each gene family of the 314G, 824G and 901G datasets. For this we inferred a ML gene trees and 100 bootstrap trees for each of the input single-gene alignments (that is, 314 × 100 bootstrap tree for the genes constituting the 314G dataset). Most gene trees did not resolve rusts, smuts and mushroom-forming fungi as monophyletic, suggesting that the phylogenetic signal of individual genes is in general not strong. Bootstrap percentages were calculated for the 3 alternative conformations for each gene and in each of the 314G, 824G and 901G dataset (Fig 1). The grouping of mushroom-forming fungi with rusts was most common among resolved trees in all three datasets, although average bootstrap percentages remain low, 33-46% for the S(R,M) as opposed to 13-19% for the R(S,M) topology. Further, on 27-48% of the bootstrap trees the rusts, smuts, agarics or combinations of these were not monophyletic. We did not find a notable correlation between the signal of individual gene trees (measured by BS%) for either the S(R,M) or R(S,M) topologies and alignment length, taxon number or evolutionary rate (results not shown).

**Figure 1.**
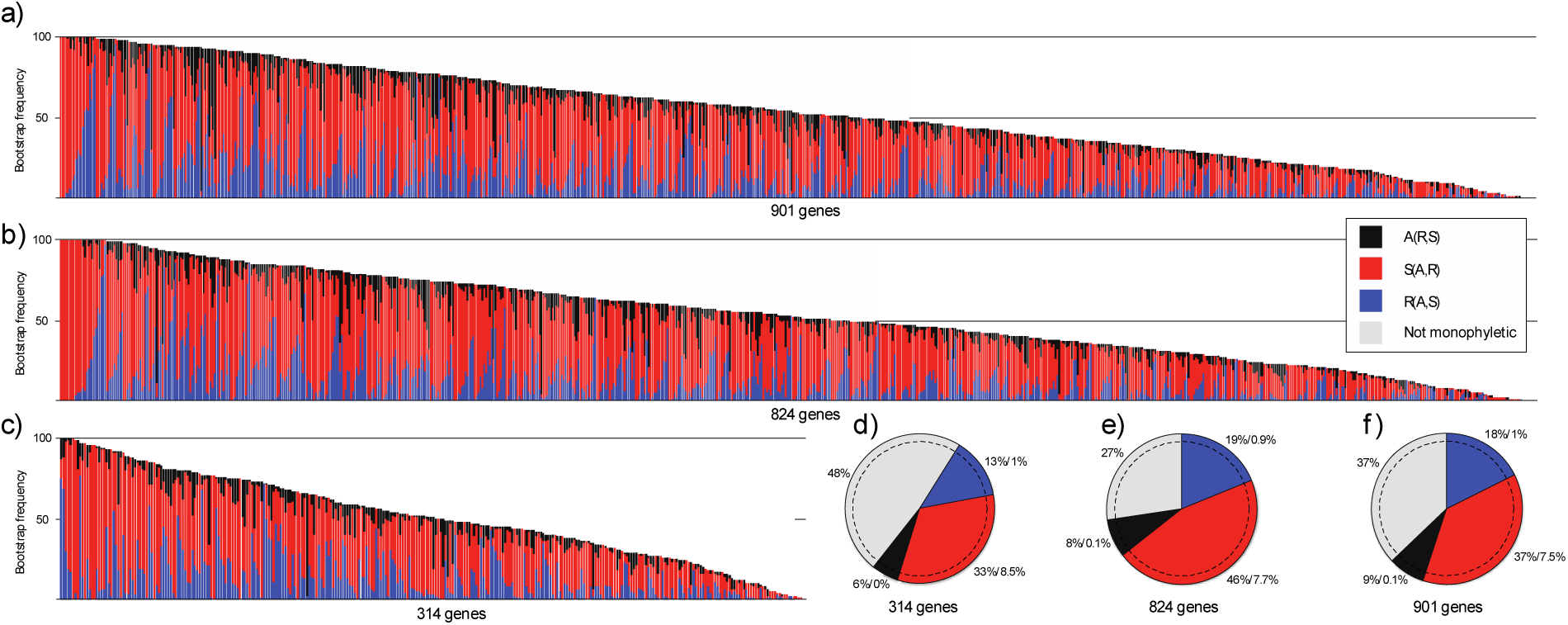
Frequencies of competing tree topologies in bootstrap samples of individual gene trees (a-c) and ML gene trees (d-f) from the 314G, 824G and 901G datasets. (a-c) the S(R,M) topology is most frequent among resolved bootstrap replicates across individual genes in all three datasets. The bootstrap frequency of the S(R,M), R(S,M) and A(R,S) topologies are shown on the y axis. Each column of the x axis corresponds to a single gene family. White section of each column represents the number of gene trees on which one or more of the large clades (rusts, smuts and mushroom-forming fungi) were not resolved as monophyletic (d-f) proportions of alternative tree topologies in ML bootstrap trees. Percentages correspond to the proportion of gene trees showing the given topology followed by the proportion of gene trees that resolved the positions of rusts, smuts and mushroom-forming fungi with >70% bootstrap support. As in the case of bootstrap trees, most genes did not resolve one or more of the large clades as monophyletic (grey section).

### Concatenated analyses

We ran partitioned ML and bootstrap analyses under best-fit models for all concatenated datasets, except the 950G dataset, which was computationally too demanding. The resulting topologies resembled each other very closely (Fig 2, Fig S3-S6), but differed in the relationships of smuts, rusts and mushroom-forming fungi. The 314G dataset placed rusts as the sister group to mushroom-forming fungi with moderate support (54%), whereas the other three datasets inferred smuts and mushroom-forming fungi as sister with the 824G and 901G datasets, providing 95% and 99% bootstrap support, respectively (Fig 3a,c). Bootstrap support for the grouping of smuts and mushroom-forming fungi showed a positive correlation with the length but an inverse relationship with the conservation of the concatenated alignments (Fig 3d).

**Figure 2.**
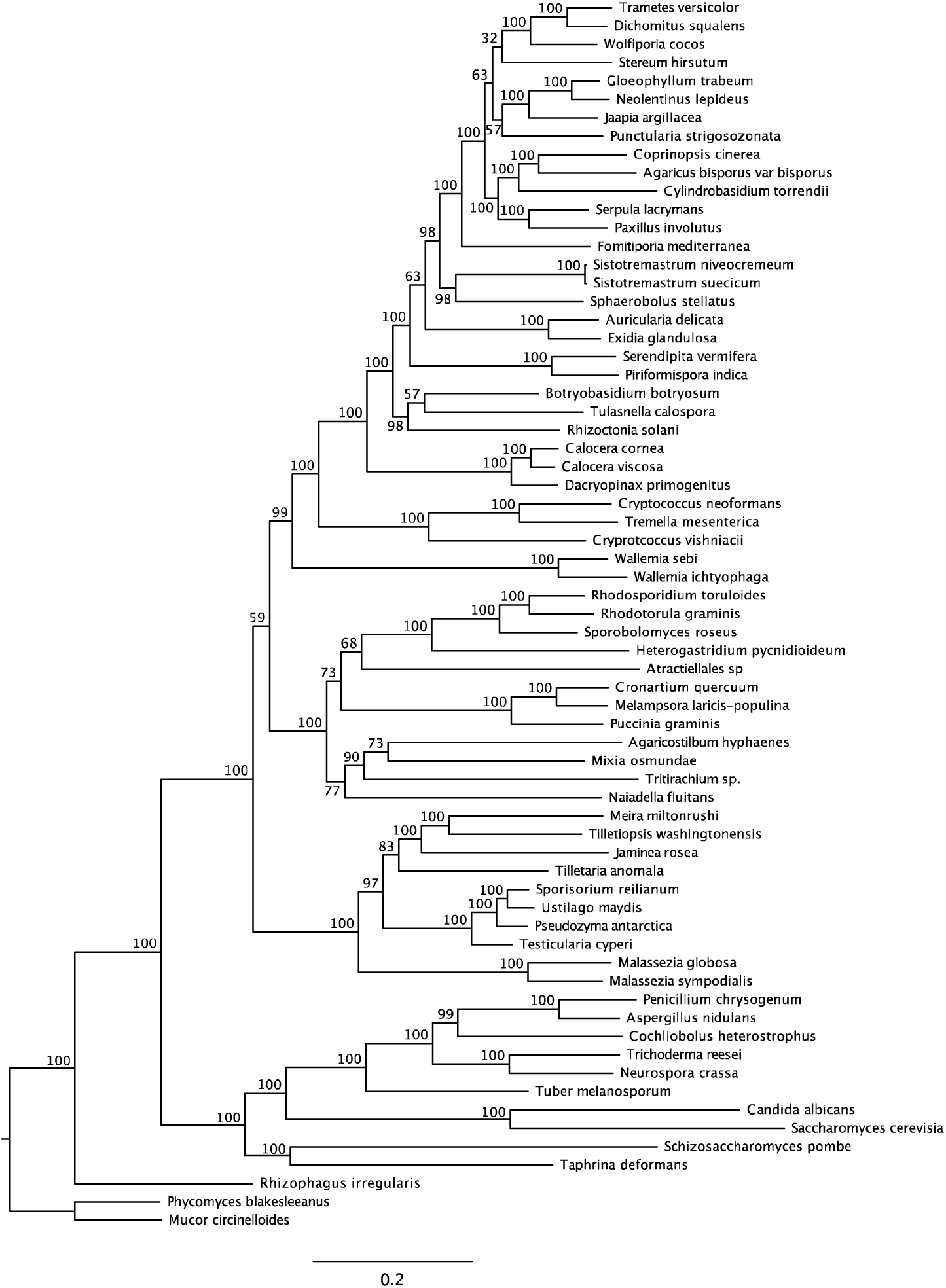
Maximum Likelihood analyses of concatenated datasets. Trees inferred from the 314G, 824G and 901G are generally strongly supported and resemble each other closely, but differ in the branching order of Agaricomycotina, Pucciniomycotina and Ustilaginomycotina. ML bootstrap percentages are shown above branches.

**Figure 3.**
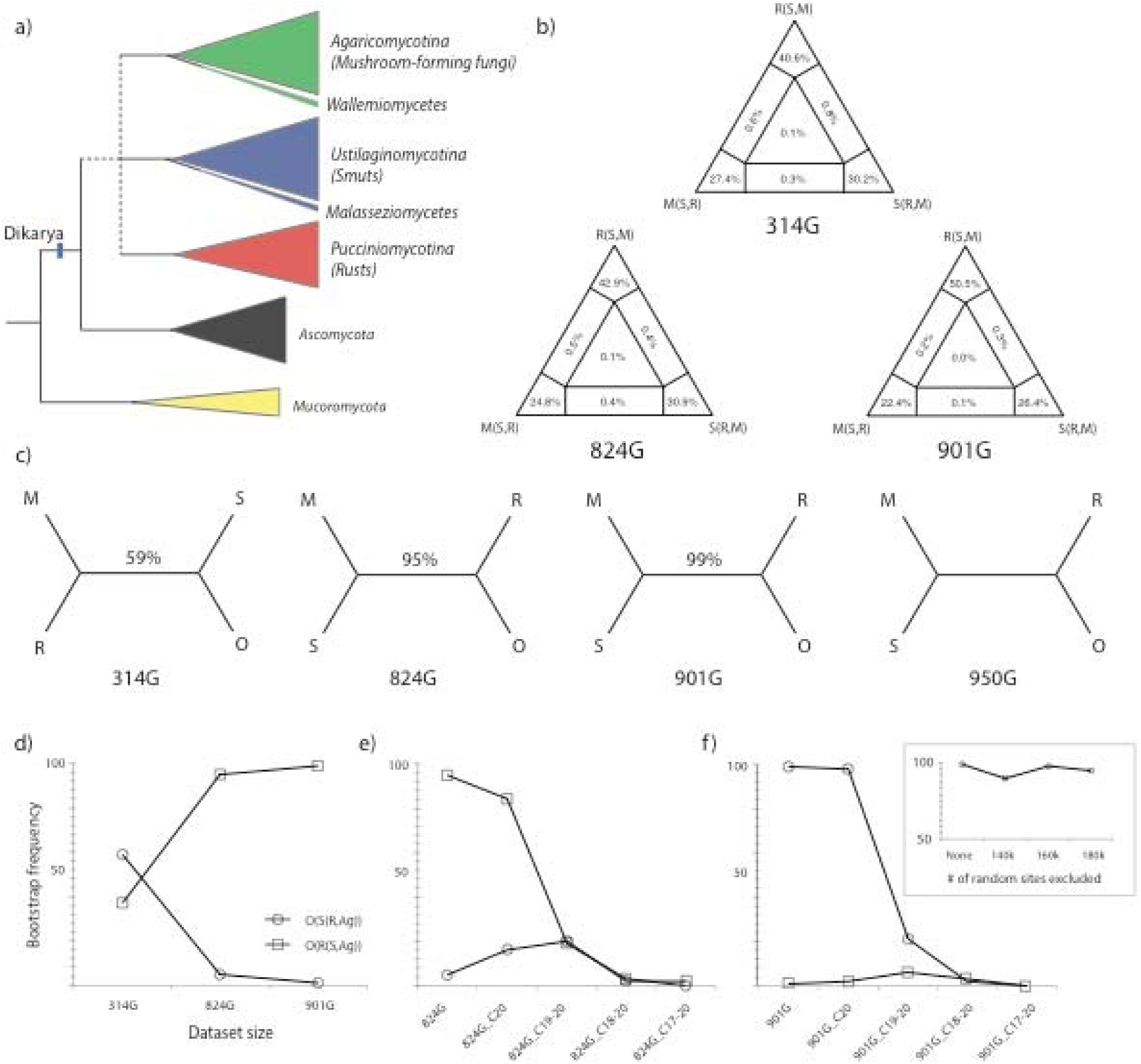
Summary of dataset properties and results of initial concatenated analyses. (a) schematic tree showing position of basal Basidiomycota splits and the Agaricomycotina (mushroom-forming fungi), Ustilaginomycotina (smuts and allies) and Pucciniomycotina (rusts and allies) in the fungal tree. The unresolved branching order of rusts, smuts and mushroom-forming fungi are shown by a dashed line. The early-diverging classes Malasseziomycetes and Wallemiomycetes are highlighted as suspected long branch taxa. (b) Results of four quartet likelihood mapping for concatenated datasets comprising 314, 824 and 901 genes referred to as 314G, 824G, 901G, respectively. (c) Simplified gene tree topologies inferred from the three above mentioned dataset plus the 950G dataset which was obtained by concatenating 950 untrimmed gene alignments. Bootstrap percentages obtained from concatenated ML analysis are shown above branches. (d) The relationship between dataset size and bootstrap proportions for smuts sister to mushroom-forming fungi (squares) and rusts sister to mushroom forming fungi (circles). These proportions change in a complementary manner when increasingly large numbers of the fastest-evolving sites are being removed from the 824G (e) and the 901G (f) datasets. Inset at (f) shows the bootstrap proprtions for the grouping of smuts with mushroom-forming fungi when equal numbers of random sites were removed from the 901G dataset.

#### The influence of dataset composition on topologies and support values

As dataset completeness and the amount of missing data can have a profound effect on inferred relationships, we examined the impact of analyzing different subsets of the original 314G dataset. First, to investigate whether the uncertainty in the topology is caused by the presence of many gene families that are indecisive with respect to basal Basidiomycota relationships, we assembled a dataset in which only decisive gene families were included. Of the original 314G dataset, only 19 gene families were indecisive with respect to rusts, smuts and mushroom-forming fungi. Restricting the analysis to gene families that had representatives in all 3 clades yielded a marginally more conserved dataset comprising 295 genes and 43,662 amino acids (295G dataset, mean pairwise genetic distance 0.404) with slightly less missing data (11.9% vs. 13% in the original 314G dataset). The tree inferred from this alignment was topologically identical with very similar support values to the tree inferred from the original 314G dataset (Fig S9), however, the bootstrap support for the node uniting rusts and mushroom-forming fungi decreased from 54% to 45%. On the other hand, concatenating only gene families that had representatives in at least 60 taxa (instead of 35) resulted in a 205 gene dataset (205G, 30,896 sites, 4.9% missing data), supporting smuts as sister to mushroom-forming fungi, with 73% bootstrap support.

We also examined what happens if we gradually remove the fastest evolving genes. By ranking genes according to their substitution rate and removing 25%, 50% and 75% fastest genes, we created three subsets of the 314G dataset: 209G (34,944 aa), 131G (23,180 aa) and 64G (11,622 aa). Analyses under the best-fit models for each partition yielded trees on which rusts grouped with mushroom-forming fungi with 61% to 71% bootstrap support.

#### Removal of fast-evolving sites

To test whether differences in support values across datasets are caused by overly variable sites, we gradually removed the fastest evolving - and potentially noisy - sites. The removal of the fastest rate categories sharply decreased bootstrap support for the grouping of smuts with mushroom-forming fungi from 99% to 98%, 21% and 0% in the 901G dataset, whereas for the 824G dataset bootstrap values decreased from 95 % to 84% and 19%. On the other hand, support for the grouping of rusts with mushroom-forming fungi increased, from 1% to 2% and 5% in the case of the 901G dataset and from 5% to 16% then 20% for the 824G dataset (Fig3/d-f, Fig S10-S21). Removal of fastest rate category did not appreciably affect the topology and support values in the 314G dataset, whereas removing the two fastest categories resulted in a topology uniting rusts with smuts (MLBS 43%, Table 1). The exclusion of further sites resulted in poorly supported topologies and the degradation of the monophyly of large groups in all three datasets, probably due to the removal of much of the phylogenetic signal.

We also specifically excluded ribosomal genes from both the 314G and 824G datasets. From the former, we excluded 29 genes (2,883 sites) whereas from the 824G dataset we excluded 63 genes (9,486 sites). Both reduced datasets yielded higher support values for the smuts as the sister of mushroom-forming fungi (Table 1). The ML topology of the reduced 314G dataset united smuts with mushroom-forming fungi as opposed to the original 314G dataset. It is noteworthy that excluding as little as 2,883 sites of ribosomal proteins was enough to flip the position of rusts and smuts on the ML topology.

#### The effects of ‘high-signal’ data subsets

Following (Shen, et al. 2017), we examined whether a small proportion of the data can dominate the inferred tree topologies. To this end, we removed from the 314G dataset the most influential amino acid sites as determined by the difference in sitewise likelihood obtained with S(R,M), S(M,S) and M(R,S) topologies. We identified the 200 sites with the largest difference in sitewise likelihoods across competing topologies for each of the 3 topologies. These sites were considered as the ones having the strongest signal for each of the 3 competing topologies. We did not observe a significant enrichment of these sites in particular genes (data not shown). The removal of top 200 strongest sites resulted in changes to tree topologies as follows. When 200 strongest sites favoring the S(R,M) topology were removed we obtained the R(S,M) topology. However, the removal of strongest sites favoring only 1 of 3 possible topologies puts the other 2 topologies at an unfair advantage, so we further removed strong sites pertaining to the other 2 alternative topologies too. Deletion of the 200 + 200 strongest sites favoring the R(S,M) and S(R,M) topologies resulted in the M(R,S) topology, whereas removal of 3× 200 strongest sites (favoring each alternative topology) yielded the S(R,M) topology, which is the same as what the original 314G dataset gave. It is notable that, consistent with (Shen, et al. 2017) analyses, the removal of small amounts of data had an impact on the topology, however, a balanced removal of strong sites did not change the original topology, after all, in our case.

Following the same logic, we examined what happens if genes with strong support for either the S(R,M) or the R(S,M) topology are analyzed separately. For this we divided the 314G dataset into ‘weak’ genes that provided <50% bootstrap for either topology and ‘strong’ genes that provided >50%. We found 65 ‘strong’ and 239 ‘weak’ genes. Interestingly, the strong genes supported the R(S,M) topology with 98% bootstrap (Fig S22), while weak genes gave the S(R,M) topology with 49% bootstrap (Fig S23). Length, rate of evolution and taxon occupancy did not show a correlation with the two sets of genes. Surprisingly, of the 65 ‘strong’ genes that conferred strong support to the R(S,M) topology when concatenated, only 10 provided strong support (>70%) to the same topology as individual genes.

#### Analyses of hard polytomies

Bayesian MCMC analyses that allow sampling polytomic trees yielded fully resolved trees for the 314G dataset under both polytomy friendly and neutral priors. These trees were consistent with the results of other unpartitioned analyses, i.e. placed smuts as the sister group of mushroom-forming fungi with strong support (BPP: 0.99, Fig S24). Because of potential differences between partitioned/unpartitioned analyses, we regard these topologies as moderately informative as to the biological question in hand, nevertheless, the analyses suggest that a hard polytomy is not a plausible hypothesis for describing basal Basidiomycete relationships. Consistent with this, FcLM analyses of the 314G dataset suggest a strong treelike pattern in the data that pertains to the split of rusts, smuts and mushroom-forming fungi, with 98.2% of the points supporting resolved topologies. The R(S,M), S(R,M) and M(R,S) topologies were supported by 40.6%, 30.2% and 27.4% of the splits, respectively. Similar patterns have been obtained for the 824G and 901G datasets (Fig 3/b). Collectively, these results suggest that the datasets contain tree-like signal with regard to the split of rusts, smuts and mushroom-forming fungi, and a hard polytomy may not plausibly explain early Basidiomycota evolution.

#### Removal of putative long branch taxa

We identified two clades sitting on long branches that could potentially be affected by long branch attraction, the Wallemiomycetes (2 species) and the Malasseziomycetes (2 spp). The deletion of one or both of these taxa from the 314G dataset had a major impact on topologies and bootstrap support (Fig 4). When we removed the Wallemiomycetes, bootstrap support for the grouping of rusts with mushroom-forming fungi increased to 82%. On the other hand, the removal of Malasseziomycetes from the same dataset gave smuts as sister to mushroom-forming fungi, supported by 73% bootstrap. Remarkably, the removal of either group yielded two of the highest bootstrap values during this study, although for conflicting topologies. Removal of both groups, on the other hand, essentially returned the original tree topology and support values (rusts + mushroom-forming fungi, 58%).

**Figure 4.**
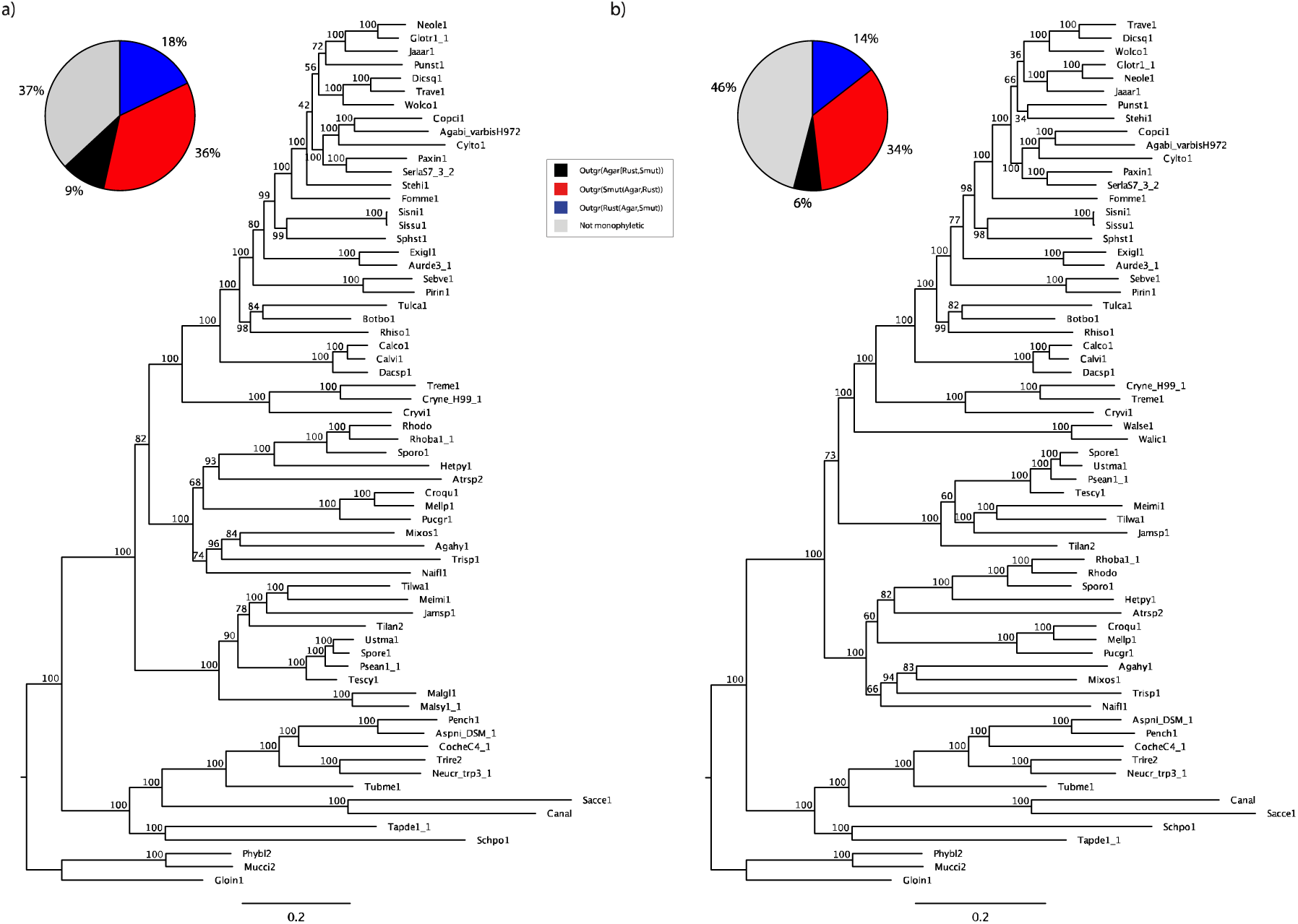
The effect of removing putative long-branch taxa. Topologies and ML bootstrap values in (a) and (b) were inferred from the 314G dataset with removing the Wallemiomycetes and the Malasseziomycetes, respectively. The distribution of individual gene tree bootstraps in the two modified datasets (pie charts) resemble closely that of the original 314G dataset.

#### Effects of model complexity

The LG+G model was selected as the best fit model for most of the partitions in all three datasets based on BIC values. In the case of the 314G dataset LG+G was preferred for 304 genes followed by RtREV+G (4 genes), MtArt+G (3 genes), WAG+G (1 gene), MtREV+G (1 gene) and CpREV+G (1 gene). We ran partitioned analyses for each dataset by specifying the best-fit model for each partition of the supermatrix and treating each input gene are a separate partition (Table 1). We then performed ML bootstrapping under modified versions of the original partitioned models (Fig 5) on the 314G dataset. First, we treated all genes as a single partition and used the LG+G model that was found to fit most individual genes the best. This was found to have a minor effect on the topology and support values. Next, we substituted the LG+G model by the WAG+G and Dayhoff+G models in partitioned, then unpartitioned analyses. Analyses under a partitioned WAG+G model yielded rusts as the sister group of mushroom-forming fungi (BS 59%) similarly to the original partitioned analysis. Under the Dayhoff+G model, however, the ML topology grouped smuts with mushroom-forming fungi (68%). Unpartitioned versions of both the WAG+G and Dayhoff+G analyses gave smuts sister to mushroom-forming fungi with 56% and 70% bootstrap, respectively. On the other hand, unpartitioned version of the analysis omitting the Wallemiomycetes or the Malasseziomycetes yielded the same topologies as their partitioned ones with increased bootstrap values.

**Figure 5.**
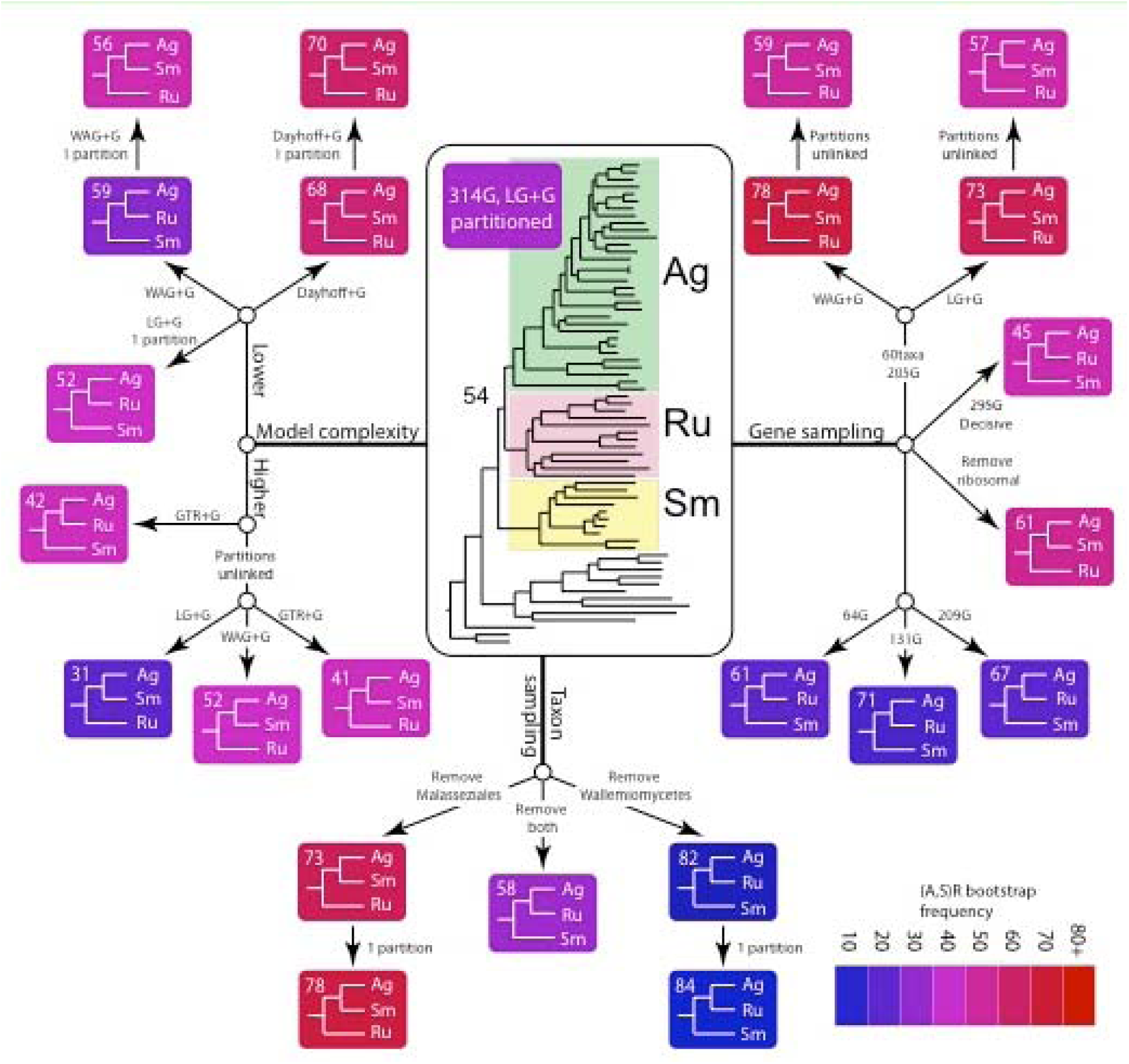
Summary of the effects of model complexity, taxon and gene sampling on topologies and bootstrap values for the relationships of rusts, smuts and mushroom-forming fungi. Boxes are colored according to the strength of bootstrap support for the grouping of smuts and mushroom-forming fungi (note that this is basically complementary to the bootstrap of rusts + mushroom forming fungi, because bootstrap trees grouping smuts with rusts had negligible frequency). Best fit partitioned models were used unless indicated otherwise.

In other experiments, we increased model complexity by unlinking model parameters across partitions. Analyses under both the LG+G and WAG+G models resulted in smuts being sister to mushroom-forming fungi, with 31% and 52% bootstrap, respectively. The more complete, 205G dataset under unlinked best-fit and WAG+G matrices across partitions grouped smuts with mushroom forming fungi, although bootstrap support decreased relative to analyses in which model parameters were linked. As a surrogate to fixed models, we employed a protein GTR+G model also. Using linked and unlinked partitioned models we obtained the S(R,M) and R(S,M) topologies with 42% and 41% bootstrap support, respectively.

### Gene tree based analyses

#### Coalescent-based species trees

As an alternative to concatenation-based methods, we examined support for basal basidiomycete relationships by using Astral-II. We estimated coalescent-based species trees from bootstrapped gene trees corresponding to input single-gene alignments of the 314G, 824G and 901G datasets. All three analyses recovered smuts as the sister group to mushroom-forming fungi, although similarly to the concatenated analyses, bootstrap support for this configuration increased with decreasing conservation of the input alignments (Fig 6). The species tree inferred from single-gene alignments of the 314G dataset had a bootstrap support of 55% whereas those of the 824G and 901G datasets supported smuts and mushroom-forming fungi as sister groups by 100%.

**Figure 6.**
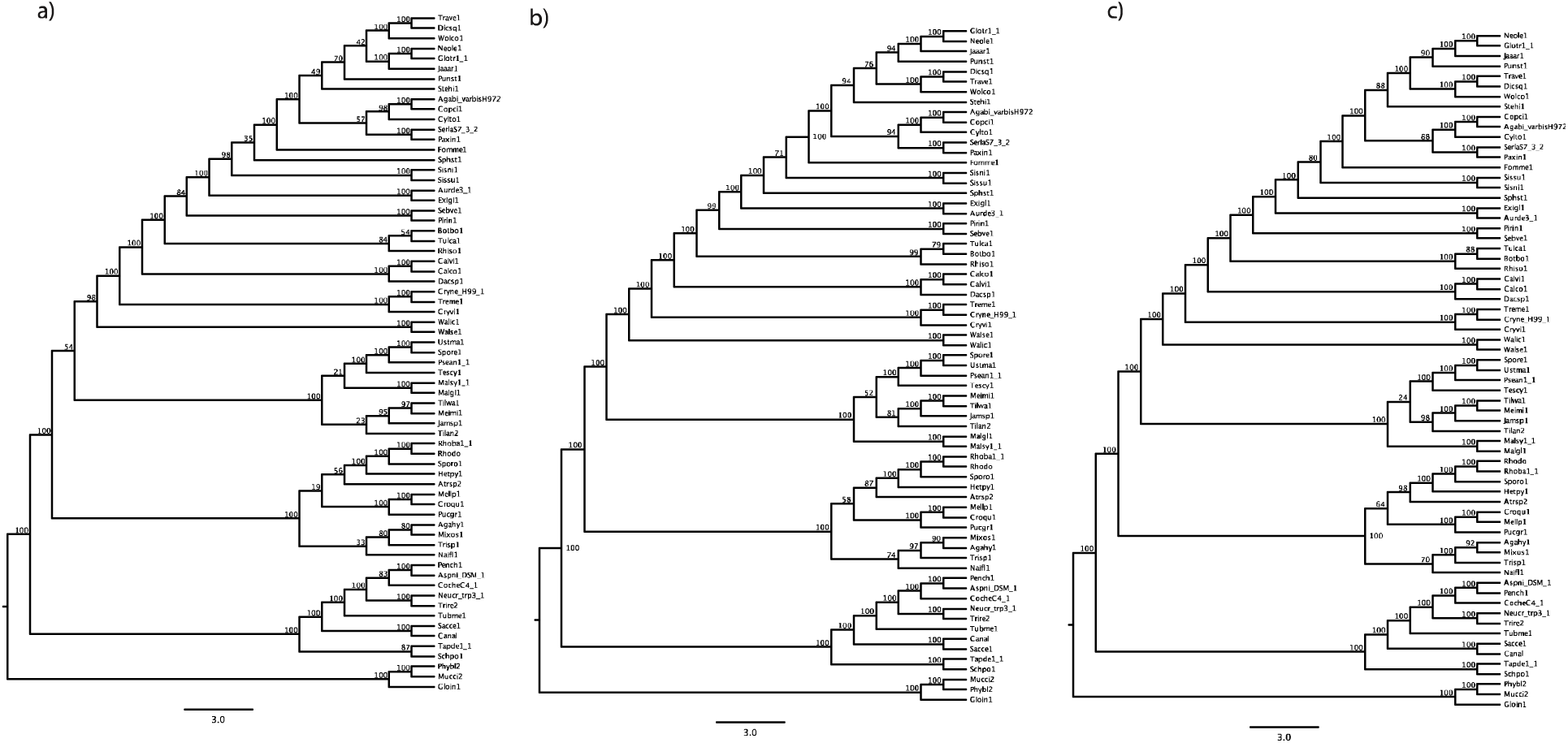
Species trees obtained using Astral-II for the 314G (a), 824G (b) and 901G (c) datasets.

### Gene content-based phylogeny

Coding gene presence/absence in Mcl-constructed clustering resulted in a binary matrix with 50,971 characters. The ML tree inferred from this dataset was very similar to the concatenation topologies and grouped rusts and mushroom-forming fungi with strong bootstrap support (98%, Fig S25). Interestingly, the Wallemiomycetes were inferred as the sister group of smuts (93%), which might be related to the reduced gene family content of both groups.

Using the reverse approach, we asked which of the competing topologies is better explained by gene presence/absence information in a parsimony context. All other branches being equal, the tree uniting rusts with mushroom forming fungi required 36 fewer steps than the tree uniting smuts and mushroom-forming fungi. Although this seems to be a marginal difference, we found it highly significant when tested in a likelihood framework (p<0.00001, approximately unbiased test).

## Discussion

### Gene tree based selection of orthogroups

Data collection is one of the primary determinants of the outcome of bioinformatic analyses. In terms of phylogenomic datasets, the distinction of orthologues and paralogues is a central task, as unwanted inclusion of paralogous genes confers incorrect topological signal and can have a large impact on inferred trees (dos Reis, et al. 2012; Dunn, et al. 2008; Whelan, et al. 2015). Our orthology detection strategy explicitly scores internal nodes of gene trees as duplications or speciations and only considers gene families not affected by deep duplications. Although this strict orthogroup selection strategy resulted in fewer orthogroups than more relaxed methods usually do (e.g. those based on best reciprocal hits), the set of gene families can be expected to virtually be devoid of paralog contamination.

### Taxon sampling and putative long branch taxa

Previous phylogenomic treatments of basal Basidiomycota involved only a single or few representatives of rusts and smuts, which were hypothesized to underlie the difficulties of resolving their relationships (Nagy, et al. 2016). In this study we increased taxon sampling density across all Basidiomycota, with special emphasis on rusts and smuts. We included whole genome sequences from 10 Ustilaginomycotina, 12 Pucciniomycotina and 32 Agaricomycotina species. Even with this improved sampling, the branching order of these major groups remained problematic. It is noteworthy, however, that we identified the Wallemiomycetes and Malasseziomycetes (Ustilaginomycotina) as potentially problematic clades, which is consistent with these taxa being problematic in rDNA and multigene phylogenies (Matheny, et al. 2007; Padamsee, et al.). Removing these resulted in the largest influence to topologies and support values, suggests they might be affected by LBA: without the Malasseziomycetes bootstrap support for R(S,M) increased to 73%, whereas without the Wallemiomycetes it increased to 82% for the S(R,M) topology. Such mutually exclusive strongly supported hypotheses are unsettling not only from a biological but also a methodological point of view. It also highlights the importance of careful taxon sampling and analyses of the contributions of individual taxa to the inferred relationships.

### Could the R(S,M) topology be an artifact of model violation?

We observed a remarkable relationship between data subsets, evolutionary models and the inferred phylogenetic relationships. First, longer and less conserved concatenated alignments favored the grouping of smuts with mushroom forming fungi, whereas more conserved subsets preferred the grouping of rusts with mushroom-forming fungi. This observation prompted us to examine whether fast evolving sites lead to smuts inferred as the sister group of mushroom forming fungi? Gradually elimination of the fastest-evolving sites decreased bootstrap support for the R(S,M) topology rapidly, whereas that of the S(R,M) topology increased slightly, until probably too many of the informative sites were deleted and support generally declined across the tree. In a complementary experiment, we created increasingly conserved versions of the 314G dataset, which gave slightly stronger support (61-71%) to the S(R,M) topology. We speculate that the stronger support by the less conserved datasets can be related to the inadequate modeling of fast-evolving aa sites in the alignment.

To address this hypothesis from the models’ side, we examined the effects of <u>model complexity</u> across a range of partitioned, unpartitioned, fixed and variable rate models as well as models with freely estimated parameters across partitions. Decreasing model complexity consistently led to increased bootstrap proportions mostly in favor of the R(S,M) topology, but also once for the grouping of rusts with mushroom-forming fungi (Wallemiomycetes removed). Analyses under unpartitioned and the poorly fitting Dayhoff model grouped smuts with mushroom-forming fungi, with up to 70% bootstrap support. On the other hand, under the most parameter-rich models (GTR, or LG+G with parameters unlinked across partitions), either smuts or rusts were inferred as the sister of mushroom-forming fungi with bootstraps between 45-55%.

These observations might indicate that poor model fit favors the R(S,M) topology whereas partitioned best-fit models and more complex models are inconclusive with regard to the branching order or rusts, smuts and mushroom-forming fungi. As in all empirical studies, the true model of evolution remains unknown and thus to what extent the data violate the assumptions of the models used here is difficult to address. Nevertheless, assessments of absolute model fit strongly rejected the best-fit models for each individual gene, suggesting that even best-fit models fail to adequately describe the process of evolution in our data. These results, combined with the results obtained using the more variable 824G and 901G datasets and by removing the fastest-evolving sites are consistent with reports of strong support for incorrect topologies arising from dataset-wide violation of model (Jeffroy, et al. 2006; Kumar, et al. 2012; Mendes and Hahn 2017; Philippe, et al. 2011; Roch and Steel 2015) assumptions. If true, however, this alerts against giving credit to the >95% bootstraps obtained for the R(S,M) topology in this and most published studies. Nevertheless, some analyses without evidently improper models also yielded smuts as the sister group of mushroom-forming fungi. For example, restricting the analysis to genes present in >60 taxa (instead of >35) yielded a 205 gene dataset with 4.9% missing data (down from 12%) resulted in the the R(S,M) topology strong bootstrap support (73-78%). Similarly, removing ribosomal genes or the Malasseziomycetes resulted in increased support for this topology.

Analyses of individual genes revealed that gene tree bootstraps most frequently supported rusts and mushroom-forming fungi as sister groups, while coalescent-based methods using Astral-II recovered smuts and mushroom-forming fungi as sisters. These results, albeit marginally, support rusts and mushroom-forming fungi as sister clades, although, it is known that under some circumstances (e.g. short internal nodes) the most common gene tree topology might differ from the true species tree, a part of the tree space known as the anomaly zone (Degnan 2013; Mendes and Hahn 2017). Similarly to concatenated analyses, (multilocus) bootstrap support increased as a function of the number of genes and their rate of evolution. This is surprising given that no such trend was observed in individual gene tree bootstraps. In general, most gene trees were poorly supported, with rusts, smuts and mushroom-forming fungi paraphyletic on >40% of the bootstrap trees. Species tree estimation from poorly estimated gene trees represent a challenging situation for gene tree-based methods (Roch and Warnow 2015), suggesting that our dataset might be challenging for Astral-based species tree inference.

### Basal Basidiomycota relationships remain an enigma

Taken together, we performed XXX analyses to understand basal Basidiomycota relationships. Our analyses uncovered multiple sources of bias that complicate phylogenomic inference in this region of the fungal tree of life, including model choice, taxon and gene sampling, or fast-evolving sites. A grouping of Ustilaginomycotina with the Agaricomycotina appears most frequently on our trees, followed by Pucciniomycotina as the sister group of Agaricomycotina, whereas Pucciniomycotina and Ustilaginomycotina as sister groups is a topology that can be rejected with confidence. Ustilaginomycotina as the sister group to the Agaricomycotina is the most common topology in both our analyses and in the literature (Ebersberger, et al. 2012; Floudas, et al. 2012; Kohler, et al. 2015; Nagy, et al. 2016; Padamsee, et al.; Sharma, et al. 2015; Toome, et al. 2013; Zajc, et al. 2013), however, it should be emphasized the we found evidence for this grouping to arise also as a result of model violation and in the presence of fast-evolving sites. This alerts against conclusively considering Ustilaginomycotina as the sister group to Agaricomycotina. On the other hand, the grouping of Pucciniomycotina with Agaricomycotina is less frequent both in our analyses and the literature and was poorly supported and found to be instable, sensitive to model choice, taxon and gene sampling.

Despite detailed examinations of the sources of incongruence, and occasional strong support for either topology, we consider basal basidiomycete relationships as unresolved. Yet, analyses rejecting the hard polytomy hypothesis suggest that there is considerable, but conflicting signal pertaining to the branching of smuts, rusts and mushroom-forming fungi. Whether and how this signal can be accessed using contemporary phylogenetic methods and models is not clear and needs further research. We speculate that the true tree topology could be restricted to a region of the parameter space that is difficult to access in the presence of patterns not captured by current models of sequence evolution. A deeper understanding of the evolutionary processes shaping fungal genomes and/or more biologically realistic models might help confidently reconstructing basal relationships in the Basidiomycota.

The relationships among rusts, smuts and mushroom-forming fungi have important implications for understanding some of the defining traits of fungi. For example, mechanisms for plugging septal pores of hyphae show great diversity among these clades, ranging from (Pucciniomycotina, Ascomycota) to pores with membrane-bounded but structurally diverse pore caps (Bauer, et al. 2006; Lutzoni, et al. 2004) in the Ustilaginomycotina and Agaricomycotina. Both the Ustilagino- and Agaricomycotina have swollen pore margins, called dolipores. If dolipores are homologous in the Ustilaginomycotina and Agaricomycotina then these two being sister groups assumes a single origin for this trait, whereas Pucciniomycotina sister to Agaricomycotina implies and extra loss. This and the morphology of spindle pole bodies (Bauer, et al. 2006) have been used as an argument for the sister group relationship between Ustilaginomycotina and Agaricomycotina. However, this hypothesis has its own challenges too. For example, septal pore anatomy can show convergent evolution (Nguyen, et al. 2017; Sharma, et al. 2015); it is not known whether the structurally different membrane caps are homologous and even losses would not be surprising given the extent of gene loss Pucciniomycotina have experienced during their evolution (Nagy, et al. 2014). Similarities between the Pucciniomycotina with the Agaricomycotina exist in a number of lesser-known traits. Many Pucciniomycotina share with Tremellomycetes (early-branching Agaricomycotina) the ability to form nanometer-fusion interaction with their hosts through tremelloid haustorial cells (Bauer, et al. 2006). This is a rare cell type developing from clamps at septa and reaching a characteristic flask-like morphology. Tremelloid haustorial cells are only found in the mycoparasitic Tremellomycetes and certain Pucciniomycotina. Mycoparasitism is further a trait shared by many Tremellomycetes and Pucciniomycotina. Further, several Pucciniomycotina orders produce gelatinous cushion-like basidiocarps (sexual fruiting bodies) similar to those of simple Tremello- and Dacrymycetes. Taken together, consistent with phylogenomic results, conflict is observed in ultrastructural and anatomical traits too; understanding how these evolved and how they can be used to distinguish between phylogenetic hypotheses for early Basidiomycota evolution will require further research.

## Conclusions

Ancient divergences in basidiomycete fungi represent a classic example of historically recalcitrant nodes (Lutzoni, et al. 2004) that pose a challenge even for genome-enabled phylogenetics (Hibbett, et al. 2013). In this study we examined sources of incongruence around basal Basidiomycota nodes in concatenation-based analyses (and, to some extent gene tree based methods). We have demonstrated that dataset composition, taxon sampling, model choice, fast-evolving sites and the analytical method all have an impact on resolving contentious relationships and that the difficulty of resolving basal Basidiomycota relationships might stem from a combination of these factors. Although we don’t have direct evidence on the direct contributions of these factors, our analyses reveal a complex interaction between the data and the analytical tools and suggest that greedy selection of sites for phylogenetic analyses can result in incorrect estimates of support values and even tree topologies. Despite identifying several sources of incongruence, however, we consider basal Basidiomycota relationships as unresolved. While this is not satisfying from the perspective of fungal biology, it highlights important challenges of genome-scale phylogenetics.

Although several of our analyses yielded apparently robust phylogenies, the pervasive conflict among trees, in our opinion, merely revealed robust incongruence rather than conclusive evidence for one or the other topology. It is becoming more and more obvious that, despite initial expectations of genome-scale datasets erasing incongruence completely from phylogenetic studies (Gee 2003; Rokas, et al. 2003), phylogenomic datasets bring about new types of challenges - that may be even more difficult to resolve than those we faced in the age of traditional phylogenetics. One such challenging situation is when individual gene trees are poorly supported, but supermatrix analyses suffer from systematic biases; in these cases both concatenation and gene tree-based analyses can perform poorly (Kubatko and Degnan 2007; Warnow 2015). We conjecture that such settings underlie the failure to resolve basal Basidiomycota relationships, creating a landscape in which the true tree topology is hidden behind complex model-data interactions. A better understanding of the evolutionary processes shaping fungal genomes and, in particular, more biologically realistic models might be needed to tackle recalcitrant phylogenetic questions either at the level of single genes or in concatenation methods.

## Acknowledgements

The authors thank Sandor Kocsube, Otto Miettinen and Gergely Szollosi for fruitful discussions on matters related to this project. This research was funded by the ‘Momentum’ program of the Hungarian Academy of Sciences (contract no. LP2014/12, by the ERC_HU_15 scheme (contract no. 118722). Genomic data used in this study were generated by the Department of Energy Joint Genome Institute and made available to the authors, which is greatly appreciated.

